# Effects of sensorineural hearing loss on formant-frequency discrimination: Measurements and models

**DOI:** 10.1101/2022.10.26.513920

**Authors:** Laurel H. Carney, David A. Cameron, Kameron B. Kinast, C. Evelyn Feld, Douglas M. Schwarz, U-Cheng Leong, Joyce M. McDonough

## Abstract

This study concerns the effect of hearing loss on discrimination of formant frequencies in vowels. In the response of the healthy ear to a harmonic sound, auditory-nerve (AN) rate functions fluctuate at the fundamental frequency, F0. Responses of inner-hair-cells (IHCs) tuned near spectral peaks are captured (or dominated) by a single harmonic, resulting in lower fluctuation depths than responses of IHCs tuned between spectral peaks. Therefore, the depth of neural fluctuations (NFs) varies along the tonotopic axis and encodes spectral peaks, including formant frequencies of vowels. This NF code is robust across a wide range of sound levels and in background noise. The NF profile is converted into a rate-place representation in the auditory midbrain, wherein neurons are sensitive to low-frequency fluctuations. The NF code is vulnerable to sensorineural hearing loss (SNHL) because capture depends upon saturation of IHCs, and thus the interaction of cochlear gain with IHC transduction. In this study, formant-frequency discrimination limens (DL_FF_s) were estimated for listeners with normal hearing or mild to moderate SNHL. The F0 was fixed at 100 Hz, and formant peaks were either aligned with harmonic frequencies or placed between harmonics. Formant peak frequencies were 600 and 2000 Hz, in the range of first and second formants of several vowels. The difficulty of the task was varied by changing formant bandwidth to modulate the contrast in the NF profile. Results were compared to predictions from model auditory-nerve and inferior colliculus (IC) neurons, with listeners’ audiograms used to individualize the AN model. Correlations between DL_FF_s, audiometric thresholds near the formant frequencies, age, and scores on the Quick speech-in-noise test are reported. SNHL had a strong effect on DL_FF_ for the second formant frequency (F2), but relatively small effect on DL_FF_ for the first formant (F1). The IC model appropriately predicted substantial threshold elevations for changes in F2 as a function of SNHL and little effect of SNHL on thresholds for changes in F1.

## INTRODUCTION

The importance of speech as a communication signal motivates studies of its neural encoding. In listeners with normal hearing, speech intelligibility is robust across a wide range of sound levels and background noise, and in the presence of temporal and spectral distortions. In contrast, listeners with even relatively small amounts of hearing loss have difficulty understanding speech in noise. A better understanding of neural speech coding would illuminate this difficulty and improve signal-processing strategies designed to aid these listeners. This study focused on perception of vowel-like sounds in listeners with normal hearing or with a range of sensorineural hearing loss (SNHL) to test the hypothesis that discrimination of spectral peaks, or formants, is consistent with a code based on the profile of neural fluctuations (NFs) in auditory-nerve (AN) responses.

In response to voiced sounds, including vowels, most AN responses are dominated by temporal phase-locking to either the fundamental frequency, F0, or to harmonic frequencies near the fiber’s characteristic frequency (CF, the frequency at which a fiber is most sensitive) (Young and Sachs, 1979; Delgutte and Kiang, 1984). The dominance of F0 wanes for fibers tuned near spectral peaks, for which responses are “captured,” or dominated, by phase-locking to the harmonic closest to CF (Deng and Geisler, 1987; Delgutte and Kiang, 1984; Young and Sachs, 1979; Zilany and Bruce, 2007). Early studies focused on the possibility that phase-locking to the temporal fine structure (TFS) of the harmonics could provide a code for spectral peaks, yet identification of neural mechanisms that decode TFS phase-locking has been elusive. The NF model focuses instead on the depth of the F0-related fluctuations in the AN responses (Carney et al., 2015; Carney et al., 2016; Carney and McDonough, 2019; Carney, 2018). These low-frequency fluctuations are important for driving responses at the level of the auditory midbrain, or inferior colliculus (IC), where nearly all neurons are sensitive to low-frequency fluctuations of their inputs (Krishna and Semple, 2000; Nelson and Carney, 2007; review: Joris et al., 2004). Most IC neurons are either suppressed or excited by a band of modulation frequencies, referred to as band-suppressed (BS) or band-enhanced (BE) neurons (Kim et al., 2020). The decoding of the NF profile is thus straightforward: average discharge rates across the population of IC neurons represent the NF profile across the population of AN fibers. These AN-response characteristics and the rate code at the level of the midbrain are illustrated in Fig. 1 for models with normal hearing (Fig. 1A) and with audiometric thresholds elevated to approximately match the mean thresholds for our group of listeners (Fig. 1B).

**Figure 1.**
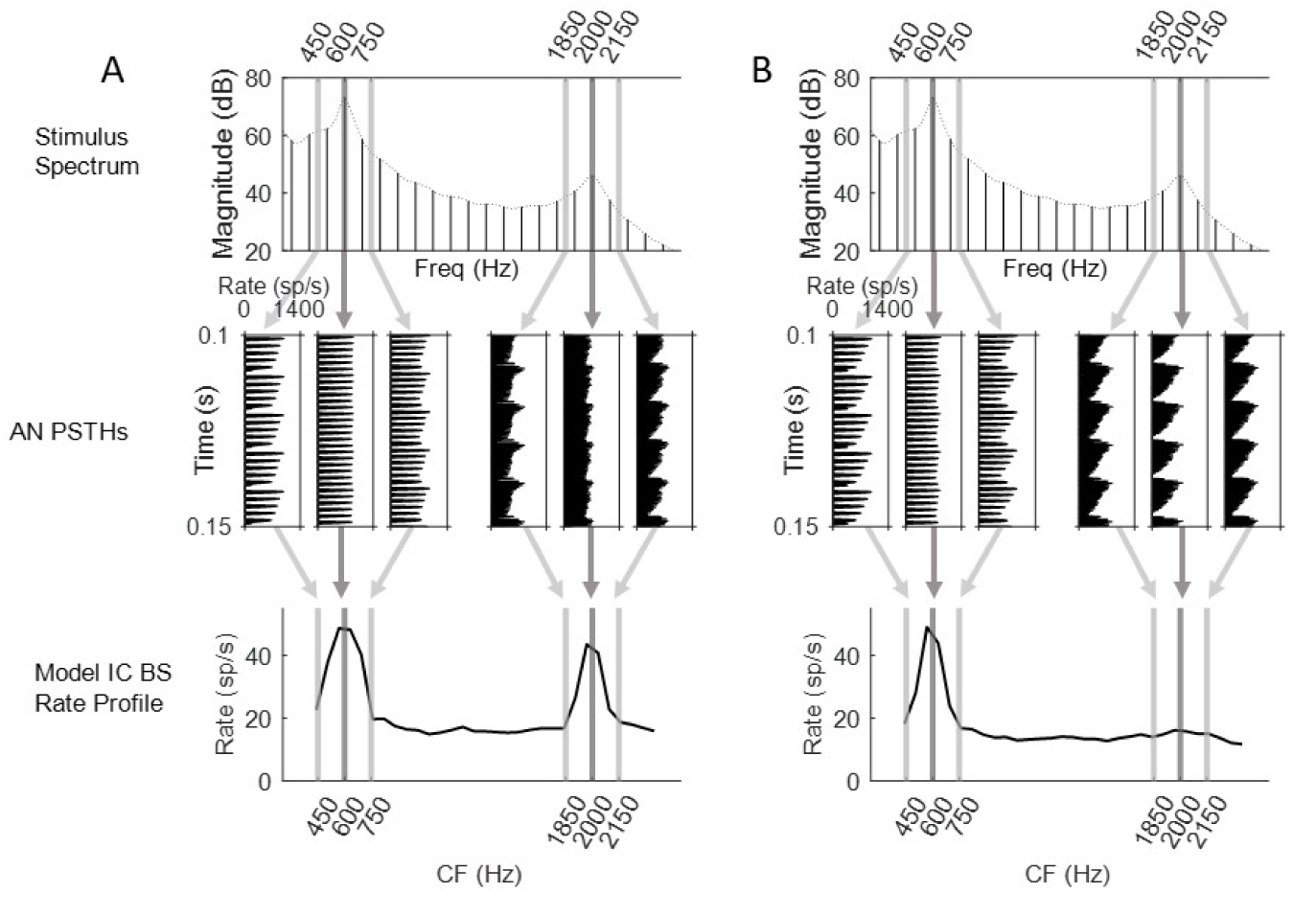
Neural fluctuations in model AN responses result in rate code for spectral peaks in the auditory midbrain. (A) Simulations of AN and IC BS responses in model with normal hearing. AN frequency channels tuned near spectral peaks have relatively shallow NFs, due to capture of IHC responses, resulting in higher rates in IC BS neurons, which are suppressed by fluctuations. (B) Simulations for model with elevated thresholds. In the impaired ear, capture of IHC responses for channels tuned near spectral peaks is weaker, resulting in larger AN NFs, and thus lower IC BS responses. Model thresholds corresponded to 12 dB HL at 600 Hz and 18 dB loss at 2000 Hz (AN model parameters: C_IHC_=0.5, C_OHC_ = 0.5). The effect of relatively mild hearing loss has a much greater impact on the encoding of F2 in model IC responses.

AN fibers tuned away from spectral peaks have large amplitude NFs at F0, due to beating between harmonics in the inner ear that is transduced by inner hair cells (IHCs). However, near spectral peaks, IHC responses saturate, and as a result the F0-related fluctuations are “flattened,” reducing the amplitude of the NFs in responses of AN fibers tuned near spectral peaks (Fig 1, middle)). The profile of NF amplitudes along the tonotopic axis thus encodes the locations of spectral peaks (Carney et al., 2015) (Fig. 1, bottom). Information about the frequencies of the spectral peaks, or formant frequencies, allow identification of vowels. Although NFs are particularly salient for voiced, or harmonic sounds, they also encode spectral shapes in unvoiced speech sounds (e.g., fricative consonants, Hamza et al., 2022). More generally, NFs have been used to successfully predict speech intelligibility in a group of listeners with SNHL (Zaar and Carney, 2022).

NFs depend on IHC sensitivity and saturation, and thus are directly influenced by cochlear gain. As a result, the reduced cochlear gain associated with SNHL affects any neural code based upon NFs. Here we tested the hypothesis that NFs encode the spectral peaks of vowel-like sounds. The bandwidths of formants were varied to increase or reduce the contrast in the NF profiles. We also estimated formant-frequency discrimination thresholds in listeners with a wide range of hearing sensitivity. We hypothesized that thresholds for all listeners would increase (worsen) as formant bandwidth was increased, and that thresholds would increase proportionally to hearing loss due to SNHL-dependent reduction in the contrast of NF profiles. Lastly, for all listeners and formant bandwidths, we varied the frequency of the formant peak with respect to the harmonic frequencies, a manipulation previously shown to affect formant-frequency discrimination (Lyzenga and Horst, 1995, 1997, 1998; Tan and Carney, 2006; Henry et al., 2017).

Our findings suggest that discrimination of F2 is quite vulnerable to hearing loss, whereas discrimination of F1 is affected very little, even by substantial hearing loss. Predicted discrimination thresholds based on the NF model were consistent with these psychophysical results. This finding has implications for the effect of hearing loss on speech understanding. The role of formant-frequency discrimination for F1 and F2 in determining the systematic patterns of vowel contrasts cross-linguistically will be discussed.

## METHODS

### Listeners

Thirty-four participants (ages 18-79 years, 26 female, 8 male) were recruited and tested using procedures approved by the University of Rochester Institutional Review Board. All participants were initially screened with a standard audiogram (0.25 – 8 kHz; Fig. 2) and were excluded if hearing loss (HL) was asymmetric (> 15 dB difference in HL between the two ears) or if HL was greater than 80 dB. Audiometric thresholds were used as input parameters for the computational models (see below), and thresholds at 500 Hz and 2 kHz were used in analyses of formant-frequency discrimination thresholds. All native English-speaking subjects (*n* = 31) were screened with the Quick Speech-in-Noise test (QSIN; Killion et al., 2004). Individual test sessions were one or two hours in duration.

**Figure 2.**
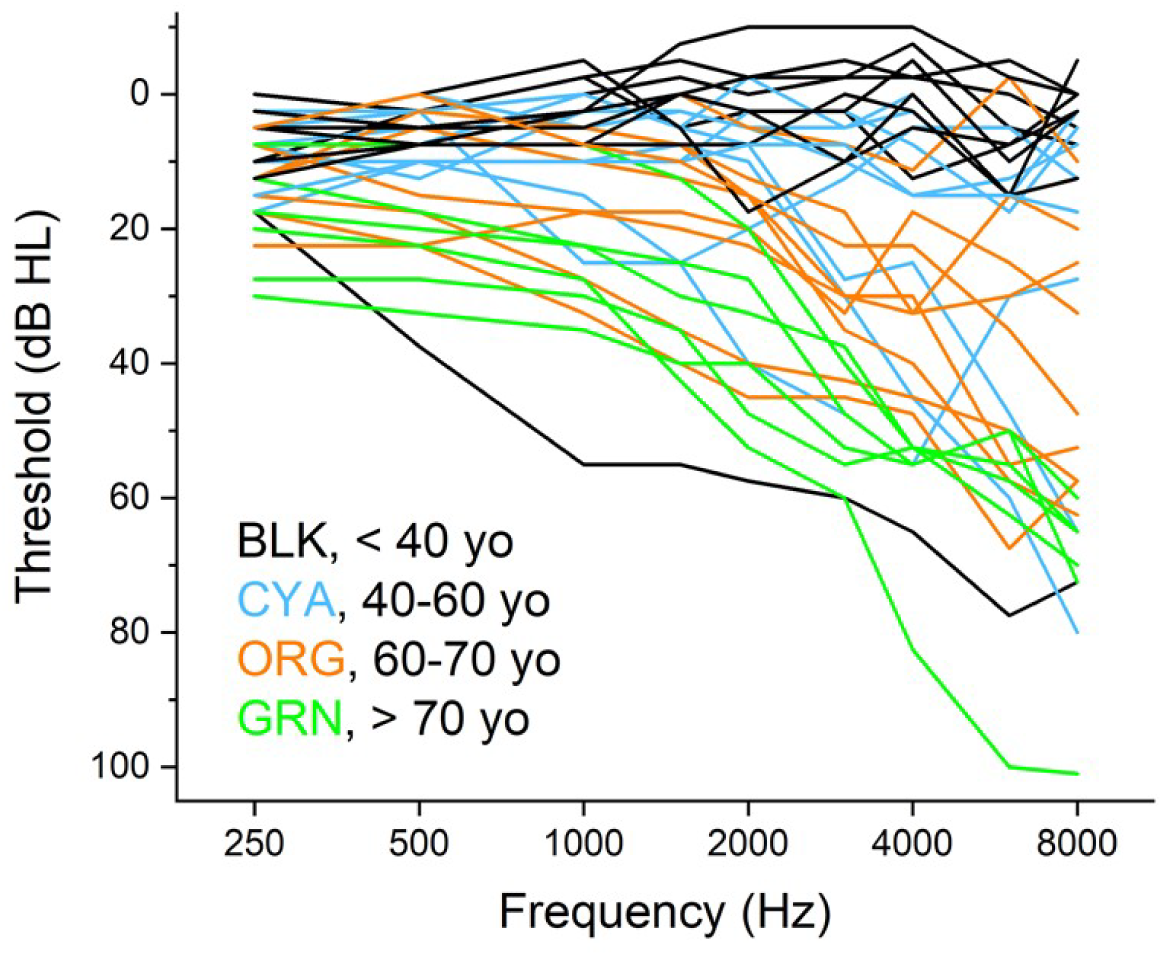
Subject audiograms. Audiograms (mean of both ears) for each subject. Age ranges are indicated by line color (see Fig. 7 for individual subject ages).

### Stimuli

Vowel-like stimuli were synthesized based on Klatt (1980), using the glottal pulse of Rosenberg (1971) and a radiation impedance implemented using a 1^st^-order highpass Butterworth filter with a corner frequency of 10 Hz. Stimuli had F0 of 100 Hz, duration of 300 ms, 25-ms cos^2^ on/off ramps, and were presented at 75 dB SPL. DL_FF_s were estimated for synthetic vowels with two formant configurations; the formants were either aligned with harmonics (designated ON; F1=600 Hz, F2 = 2000 Hz) or midway between harmonics (designated BTW; F1 = 650 Hz, F2 = 2050 Hz) (Fig. 3). DL_FF_s were also estimated for three formant bandwidths (50, 100, and 200 Hz; Fig. 3). The 12 stimulus conditions were tested in blocks that were randomly ordered for each subject.

**Figure 3.**
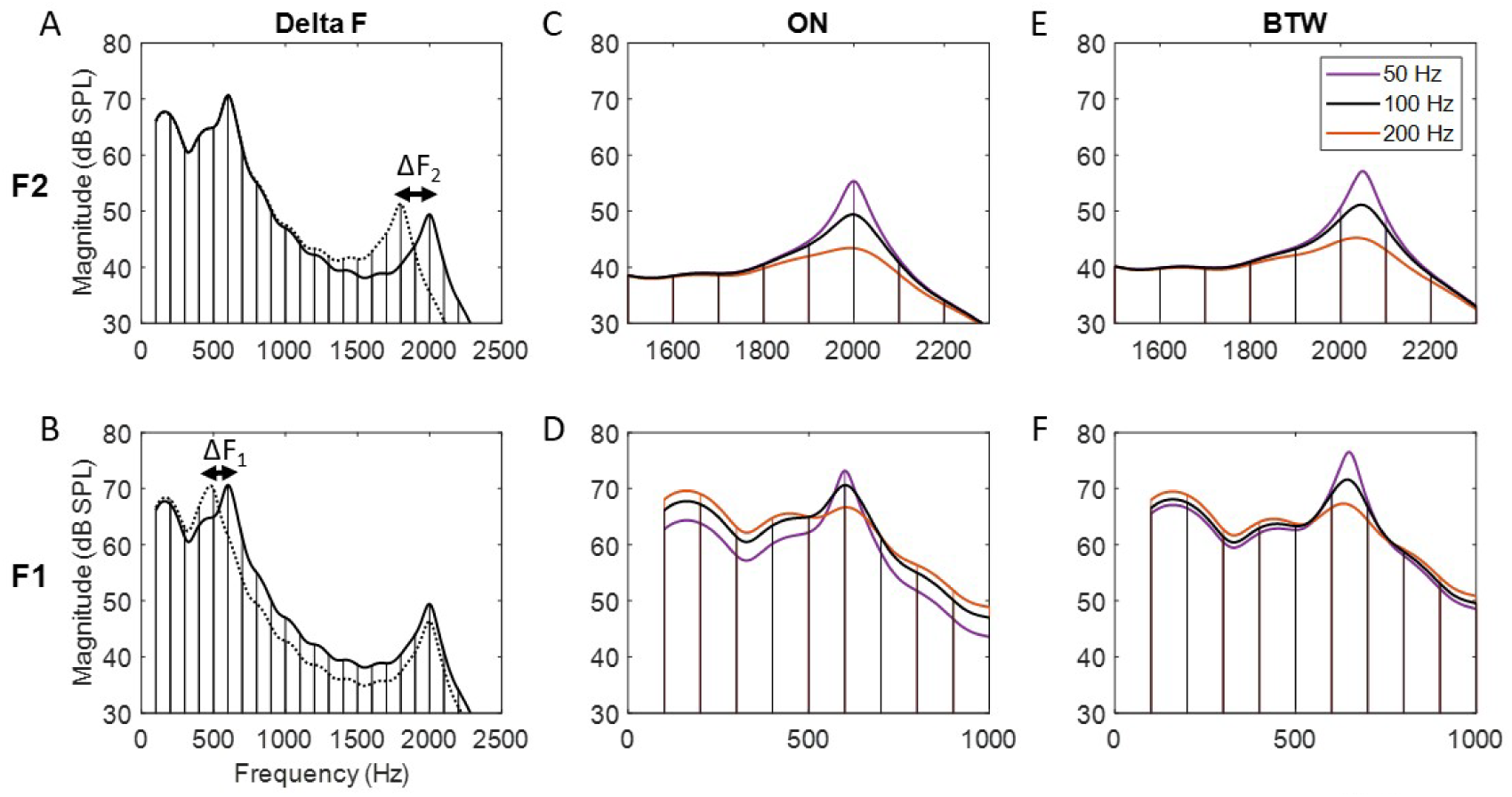
Stimulus spectra. Discrimination thresholds were measured using a standard pair of identical sounds and a target pair with a change in (A) F2 or (B) F1. Spectral peaks were either centered (C, D) on (ON) a harmonic, or (E, F) between (BTW) harmonics. Formant bandwidths were 50 (purple), 100 (black), or 200 (orange) Hz.

### Procedure

A two-interval, forced choice, two-down one-up tracking procedure was used, as follows. Each trial contained two pairs of sounds; the standard pair had identical sounds, and the target pair had sounds that differed in either F1 or F2. For each track the formant-frequency difference (ΔF) started at 40%. The formant-frequency difference was reduced by a factor of two following every two correct responses, or increased after incorrect responses, until two reversals occurred. ΔF was then reduced or increased by a factor of 1.26 until four reversals occurred, then reduced or increased by a factor of 1.12 until a total of ten track reversals were attained. Individual tracks were retained for analysis if a criterion of (standard deviation of ΔF / track threshold) < 0.3 was achieved. The DL_FF_ for each 10-reversal track was estimated as the average ΔF of the final six reversals. Overall threshold for a particular stimulus condition was based on the average of at least three individual track thresholds. If the ratio of the standard deviation of the threshold estimates to the mean threshold was > 0.3, additional tracks were collected until the criterion (< 0.3) was achieved. Statistical analyses were based on ANOVA analyses of linear mixed-effects models, focusing on the bandwidth factor and its interaction with the factors of HL, QSIN score, and Age.

### Computational Modeling Methods

Model thresholds were estimated as follows: auditory-nerve (AN) responses were simulated for 11 high-spontaneous-rate fibers with log-spaced CFs spanning 1/3 octave, centered at the formant frequency being tested. A single model AN fiber was simulated for each frequency channel. Thresholds based on individual audiograms for each listener were included in the AN model (Zilany et al., 2014).

Model DL_FF_s were based on Band-Suppressed (BS) IC model neurons (Fig. 4; Carney and McDonough, 2019). A single model IC neuron was simulated for each of the 11 frequency channels set up by the AN model, spanning 1/3 octave above and below the formant frequency being tested. Midbrain neurons were either excited or suppressed by temporal fluctuations on their inputs. The BS population rate-profile peaks at frequencies near formants, where fluctuations are reduced by capture (Fig. 1). Fluctuations were minimal near formant peaks, due to capture of the IHC responses by a single harmonic near CF (Deng and Geisler, 1987; Zilany and Bruce, 2007). BS models had modulation transfer functions with a trough frequency at 64 Hz, the median observed in a population of rabbit IC neurons (Kim et al., 2020).

**Figure 4.**
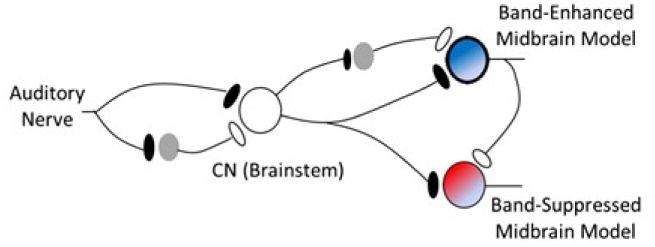
Schematic illustration of models for band-enhanced (BE) and band-suppressed (BS) IC neurons. Model IC BS neurons were used in this study. (From Carney et al., 2015)

The stimulus interval with model population rate-profiles having the largest mean squared difference across the pair of stimuli was chosen as the target interval. Rate profiles were based on the mean rates of model IC BS neurons, including both onset and sustained responses. Although BE model neurons could have been included in threshold predictions, the two model response types are closely related (Fig. 4), thus the benefit of including both types would have been a straightforward enhancement of sensitivity. There is not yet sufficient evidence concerning the projection patterns of neurons with BS vs. BE MTFs, or how these two apparently opponent response types might be combined in the central nervous system (Kim et al., 2020). Although inclusion of both response types could potentially improve discrimination sensitivity, for simplicity, we opted to base the threshold predictions reported here on the population response of BS cells.

Thirty trials were run for each ΔF in every stimulus condition, i.e., for variation of F1 or F2, for standard stimuli with formants on or between harmonics, and for the three formant bandwidths. As in the psychophysical tests, each trial consisted of two pairs of stimuli: the standard interval pair contained two identical stimuli with the formant at the standard frequency; the test pair contained one interval with the standard formant frequency, and the other with a shifted formant frequency. Half of the simulated trials had the test stimulus as the first interval of the test pair, and half had this stimulus as the second interval, in order to counterbalance any effects of neural adaptation in the model. The stimulus pair with the larger mean-squared error between neural population rate profiles to the two intervals in the pair was selected as the target interval. Percent-correct model performance was based on the number of trials for which the test pair was correctly identified. Simulations were performed for %ΔF = 0.01, 0.02, 0.05, 0.1, 0.2, .5, 1, 2, 5, 10, 20, and 50 to cover a wide range of thresholds. For listeners with audiometric thresholds greater than 30 dB HL, the range of %ΔF2 was further extended to 95% Each DL_FF_ estimate was based on a logistic function fitted to %-Correct vs. %ΔF. Model threshold was estimated as the %ΔF that resulted in 70.7% correct performance. Data files and Matlab code used for simulations and figure preparation are available at https://osf.io/tnjup/.

## RESULTS

### Thresholds vs. HL at formant frequencies

Subject thresholds for the F2 and F1 conditions, as a function of HL and stimulus bandwidth, are presented in Fig. 5; results from ANOVA analysis of linear mixed-effects models for the same data are presented in Table 1. For the F2 condition there was a strong, positive, linear correlation between threshold and HL. These correlations were similar for all three tested bandwidths, and also for both ON and BTW stimuli (see Fig. 5 legend for linear regression R^2^ values). A significant bandwidth-dependent increase in threshold was also observed for both ON and BTW, being particularly evident for subjects with HL < 15 dB (*cf*. Fig. 5 top row panels), and confirmed with ANOVA analysis (Table 1). For the F1 condition there was little evidence for positive correlation between threshold and HL for any of the tested bandwidths (Fig. 5, bottom row; Table 1). Similar to the F2 condition, a significant bandwidth-dependent increase in threshold was observed in the F1 condition for both ON and BTW, particularly evident for subjects with HL <15 dB, and was confirmed by ANOVA analysis (Table 1). These results are consistent with a statistically significant, positive, bandwidth- and mid-to-high frequency HL-dependent difficulty in discriminating differences in F2, a difficulty that is comparatively absent for F1.

**Figure 5.**
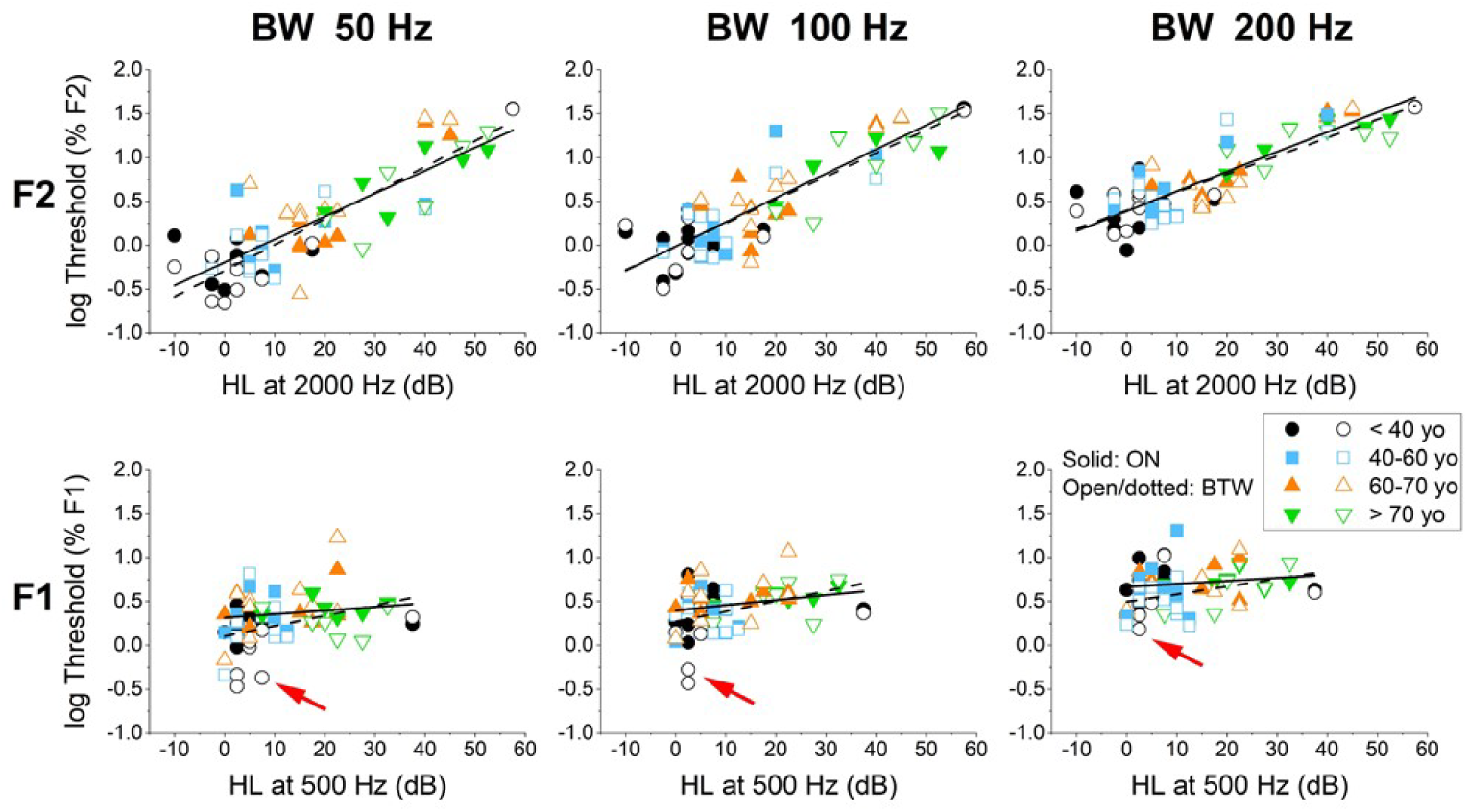
F2 and F1 threshold as a function of HL and stimulus bandwidth. For each plot the aggregate data for ON (solid symbols) and BTW (open symbols) are fitted with linear regressions (solid and dotted lines, respectively); data points are color- and symbol-type-coded by age range as indicated (see also Fig. 7), plotted as log_10_(%F). For the F2 condition (top row), thresholds for both ON and BTW stimuli increased significantly with increasing HL at 2000 Hz (linear regression R^2^ values: ON, 0.72 (50 Hz), 0.74 (100 Hz), 0.75 (200 Hz); BTW, 0.70 (50 Hz), 0.72 (100 Hz), 0.71 (200 Hz)). For the F1 condition (bottom row) there was little evidence for correlation between threshold and HL at 500 Hz (linear regression R^2^ values: ON, 0.02 (50 Hz), 0.06 (100 Hz), 0.01 (200 Hz); BTW, 0.08 (50 Hz), 0.11 (100 Hz), 0.11 (200 Hz)). In both the F2 and F1 conditions, for HL < 10 dB discrimination thresholds were elevated for both ON and BTW for BW 200 Hz compared to the narrower bandwidths. See Table 1 for ANOVA analyses of the plotted data. Note the age-dependence of BTW thresholds for F1 discrimination, as indicated in Table 1, is highlighted by the red arrows, which point to the relatively low thresholds for F1 BTW for younger listeners with normal hearing (NH).

**Table 1.** P-values of ANOVA analysis of Linear Mixed-Effects Models, focusing on the bandwidth factor and its interaction with the factors of hearing loss (HL), QSIN score, and age. The * symbol indicates statistical significance (p < 0.05); the † symbol highlights the age-dependence of threshold for the F1 BTW condition (see arrows in Fig. 5).

| **Factors and Interactions** | **F1 ON** | **F1 BTW** | **F2 ON** | **F2 BTW** |
| --- | --- | --- | --- | --- |
| Bandwidth | p < 0.001* | p < 0.001* | p < 0.001* | p < 0.001* |
| HL @ 500 (F1) or<br>2000 (F2) (N = 34) | 0.20 | 0.02* | p < 0.001* | p < 0.001* |
| Bandwidth | p < 0.001* | p < 0.001* | p < 0.001* | p < 0.001* |
| HL:Bandwidth | 0.68 | 0.75 | 0.08 | 0.004* |
| QSIN scores (N = 31) | 0.48 | 0.18 | 0.013* | 0.009* |
| Bandwidth | p < 0.001* | p < 0.001* | p < 0.001* | p < 0.001* |
| QSIN:Bandwidth | 0.16 | 0.34 | 0.64 | 0.015* |
| Age (N=34) | 0.14 | 0.007*† | 0.001* | p<0.001 |
| Bandwidth | p < 0.001* | p < 0.001* | p < 0.001* | p<0.001 |
| Age:Bandwidth | 0.47 | 0.02* | 0.20 | 0.02 |

Ratiometric comparison of ON and BTW thresholds as a function of HL and stimulus bandwidth are presented in Fig. 6. For subjects with HL less than 10 dB at either 2000 or 500 Hz, thresholds for ON were typically higher than BTW for both the F2 and F1 conditions, particularly for subjects with age less than 40 yrs (black symbols, Fig. 6). This trend was reduced for all subjects, and for both the F2 and F1 conditions, as a function of increasing stimulus bandwidth. These results indicate that most subjects had higher thresholds for the ON condition (i.e., threshold ratios > 1) as compared to the BTW condition, for both the F2 and F1.

**Figure 6.**
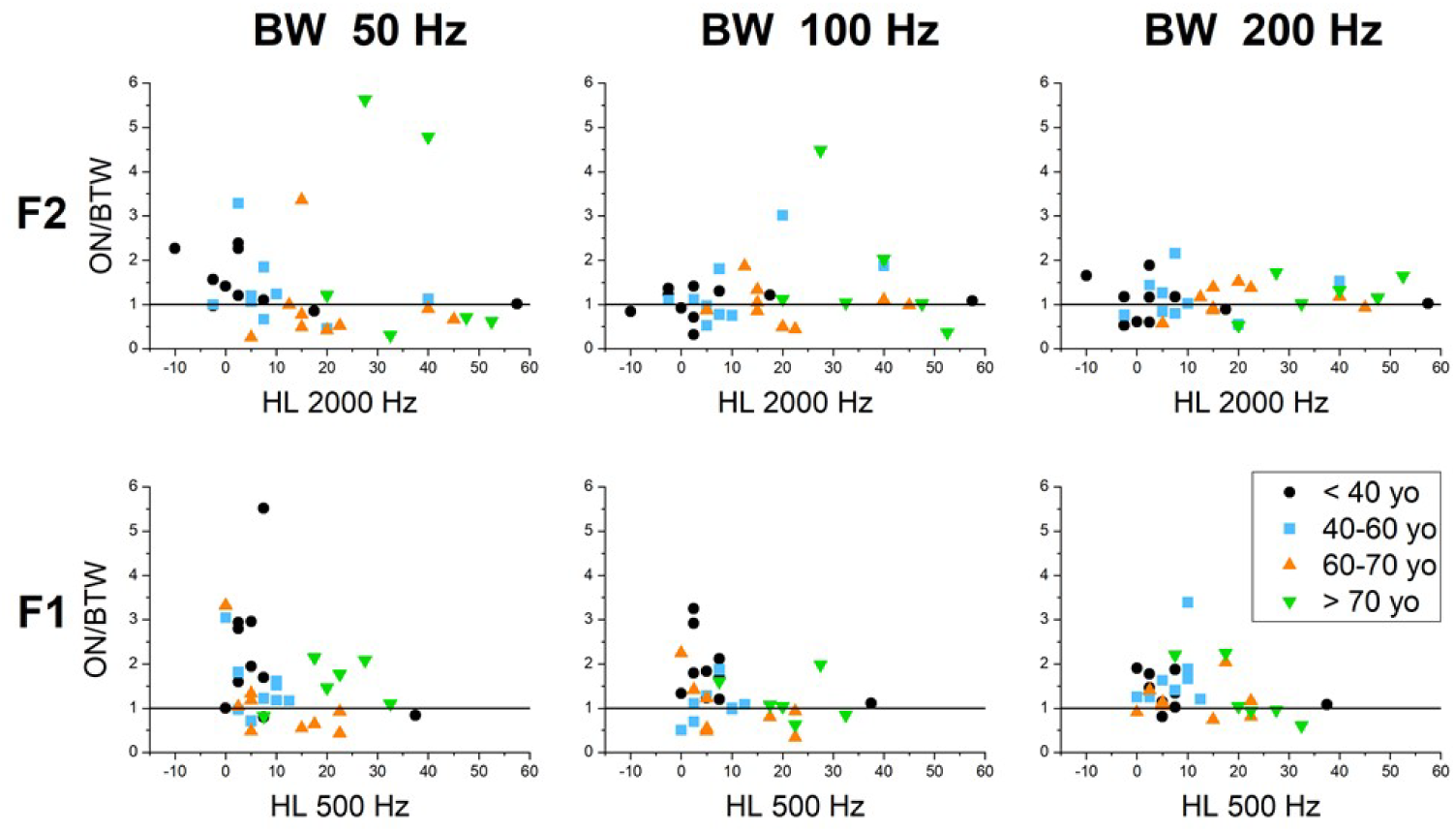
Ratio of ON and BTW thresholds as a function of HL and bandwidth. For F2 (top row), normal-hearing subjects (<15 dB HL) tend to have greater thresholds for ON than BTW at 50 Hz bandwidth, a trend that is absent at greater bandwidths; subjects with mild-to-moderate HL tend to have greater threshold for ON compared to BTW as a function of increasing bandwidth. For F1 (bottom row), normal-hearing subjects have a consistently higher threshold for ON than BTW, independent of HL and bandwidth; the thresholds for ON and BTW are similar with increases in HL and bandwidth.

QSIN scores as a function of subject age and HL are presented in Fig. 7. The strongest linear correlation (i.e., regression R^2^ value) was observed for HL at 2000 Hz (Fig. 7, right panel). This result was supported by ANOVA analysis (Table 1). Outlier QSIN scores > 10 for two subjects suppressed these correlations, particularly with respect to HL; when those two data points were excluded the R^2^ values were 0.17 (age), 0.38 (HL 500 Hz), and 0.50 (HL 2000 Hz). These results indicate a general, positive correlation between QSIN score and HL, and further indicate that formant-frequency discrimination tasks, such as those used in this study, could be used as non-cognitive surrogates to the QSIN test to assess HL and vowel-sound discrimination.

**Figure 7.**
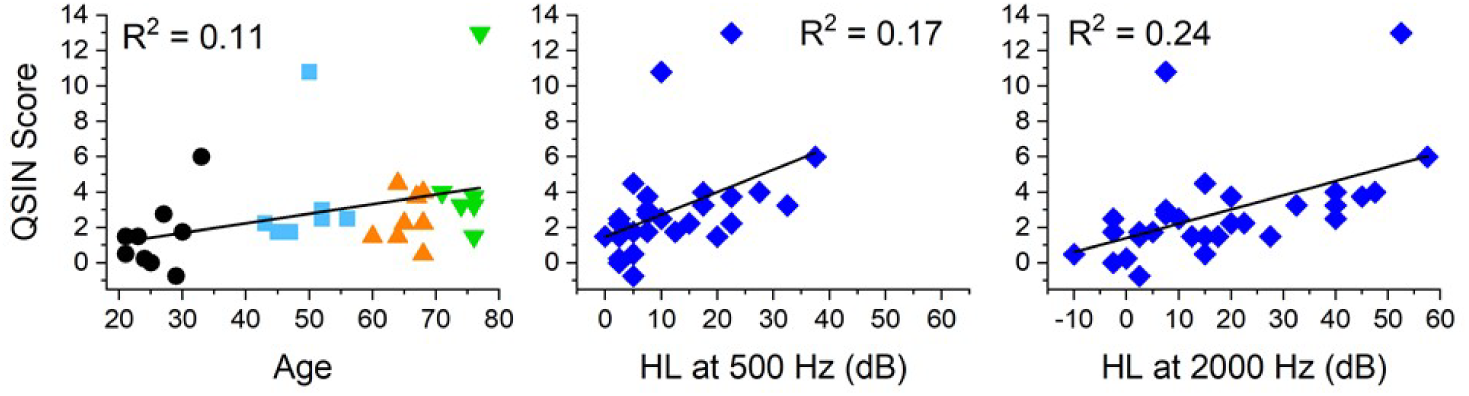
QSIN scores as a function of age and HL. Linear regressions and R^2^ values are indicated; color and symbol-type (left panel) match that of Figs. 5 and 6. QSIN scores were correlated with age and HL, particularly for HL at 2000 Hz (right panel). See Table 1 for ANOVA analyses of these data.

### Statistical Analysis

Statistical evaluation (*p*-values of ANOVA analysis of linear mixed-effects models) of the human-derived thresholds is summarized in Table 1. For both F1 and F2, and in both the ON and BTW conditions, bandwidth had a significant effect upon DL_FF_, with no significant interaction between bandwidth and age. There was a significant effect of age (green box, Table 1) and hearing loss upon the F1 BTW condition, for which some young subjects with low audiometric thresholds had especially low thresholds (presented graphically below). For F2, but not F1, QSIN score, age, and hearing loss at 2 kHz all had significant effects on DL_FF_s for both the ON and BTW conditions. Interactions between bandwidth and hearing loss, and between bandwidth and QSIN score, were significant only for the F2 BTW condition. The effects of hearing loss near the formant frequency, QSIN score, and age were always significant for the F2 thresholds, the only significant interaction for the F1 BTW condition being that between age and bandwidth. These statistical analyses indicated that all interactions between a factor (age, QSIN score, and hearing loss) and bandwidth were significant for the F2 BTW condition.

Listeners’ formant-frequency discrimination thresholds were estimated based on AN and IC BS computational-model responses. Predictions based on AN rate profiles quantify the ability to predict responses based on AN rate-based coding of differences in the stimulus spectra. These results can be compared to the predictions presented below that were based on model IC BS neurons, which quantify the ability of neural fluctuation-based coding of the spectra.

The rate profiles for 1/3-octave CF ranges of AN fibers are shown in Fig. 8, for stimuli that differed in F1 (bottom row) or F2 (top row) conditions, ON (on-peak) or BTW (between-peak) conditions, and formant bandwidth (colors). Responses to the standard stimuli, with spectral peaks at 600 or 2000 Hz for the ON condition, or 550 or 1950 Hz, for the BTW condition, are shown in dark lines, with line color indicating formant bandwidth. Responses to target stimuli that were shifted down in frequency by 5% of the formant frequency are shown in light lines, for all three bandwidth conditions. Stimuli were presented to the model in two pairs, one with identical standard stimuli, and one with a formant-frequency difference. The sum of the squared differences across the 11 CF channels was used to estimate the stimulus pair that differed on each trial. Percent-correct scores were tallied across 30 trials for each stimulus condition, for a range of formant-frequency differences. A logistic function fit to the percent-correct results was used to estimate the formant-frequency differences corresponding to threshold, or 70.7% correct.

**Figure 8.**
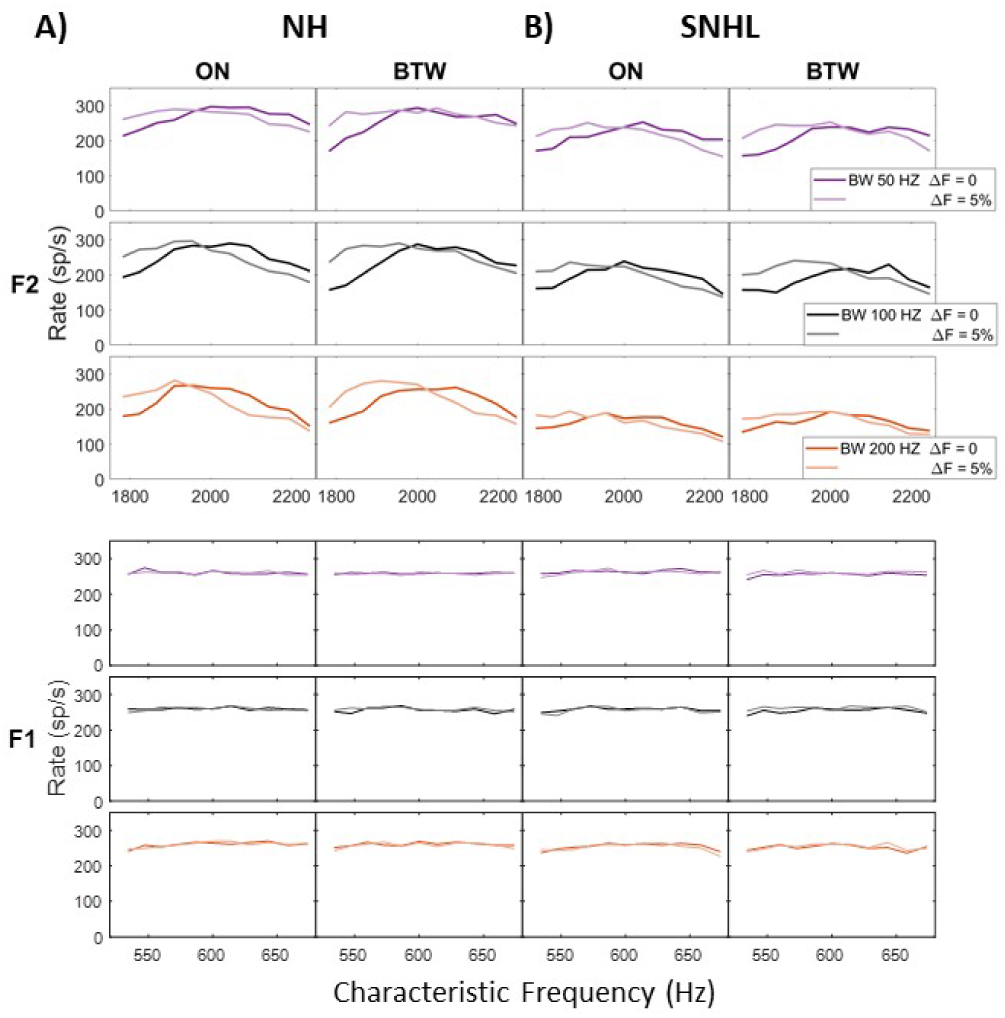
AN model rate profiles for models with (A) NH and (B) SNHL corresponding to 12 dB loss at 600 Hz and 18 dB loss at 2000 Hz (AN model parameters: C_IHC_=0.5, C_OHC_ = 0.5). AN model rate profiles are illustrated for stimuli that differed by ΔF = 5%, which is above threshold for either NH or listeners with the degree of SNHL included in (B) (cf., Fig. 5, note log_10_(5%) ≈ 0.7). Model rate profiles are shown for the three formant bandwidths using line colors matched to Fig. 3: 50-Hz BW, purple; 100-Hz, black; 200-Hz, orange. For each BW, the darker line corresponds to the standard stimulus, and the lighter line is the response to a formant shifted down by 5% of the standard formant frequency (2000 Hz or 600 Hz). Each AN model profile is for 11 frequency channels spanning 1/3-octave centered at the formant frequency. Model threshold estimates were based on the sum of squared errors between rate profiles. The rate profiles illustrated here are based on averages of 10 trials.

The AN-model threshold estimates as a function of subject thresholds are presented in Fig. 9. In the F2 condition, for bandwidths of 100 and 200 Hz, the model-derived thresholds were positively correlated with subject thresholds and roughly captured the human threshold values (R^2^ range of 0.28-0.36 for ON and BTW). For the F1 condition, for all three bandwidths, the AN model consistently returned elevated thresholds (points above the unity line on Fig. 9) that were negatively correlated with the human threshold values plotted as a function of SNHL. These model results suggest that the AN model captures some aspects of human F2 and F1 thresholds, but that absolute AN-model threshold values were consistently higher than human thresholds for the F1 conditions and for the 50-Hz F2 condition. The relatively small AN rate differences observed in the F1 condition responses, for both NH and SNHL models (Fig. 8A, B, bottom row) are consistent with the elevated threshold estimates in Fig. 9. Differences in AN rate profiles for stimuli that differed in F2 (Fig. 8A, B, top row) were large enough to estimate thresholds similar to some listeners’ thresholds, but the trends with SNHL and BW in the listeners’ threshold were not well predicted by AN model responses.

**Figure 9.**
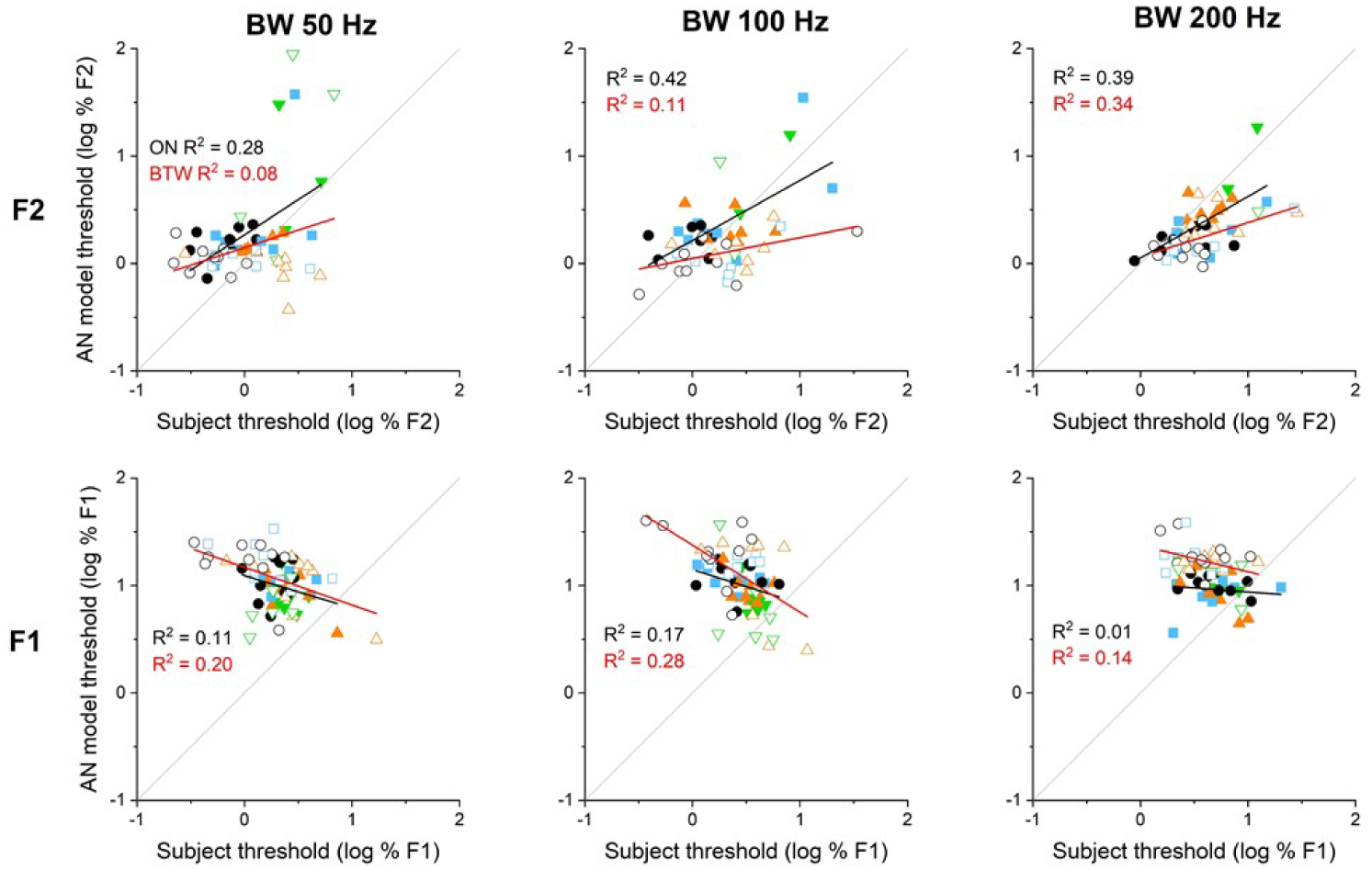
AN-model thresholds as a function of subject thresholds. Symbol types and colors as in Fig. 5, with the gray line indicating unity. Bold lines correspond to linear regressions of the displayed ON (black) and BTW (red) data sets, with the R^2^ values indicated in each panel. The AN model captured some of the F2 data trends for 100 and 200 Hz bandwidths, but less so for the 50-Hz bandwidth (cf. the respective panels in Fig. 5). For F1 conditions, AN-model thresholds were consistently elevated with respect to human thresholds (points above the unity line) and were negatively correlated with the listeners’ thresholds. Model thresholds were based on 30 repetitions of single-trial responses that were used to estimate logistic functions (see Methods).

Figure 10 shows examples of IC BS model rate profiles for a NH model (Fig. 10A) and a model for listeners with elevated audiometric thresholds (Fig. 10B; audiometric thresholds are matched to those illustrated in Fig. 1). The AN inputs used for the example IC BS model responses shown in Fig. 10 were the exact responses shown in Fig. 8. Thus, direct comparison of Figs. 8 and 10 provides a qualitative illustration of the difference in information between AN rate profiles and AN NF profiles, which are mapped into IC rate profiles.

**Figure 10.**
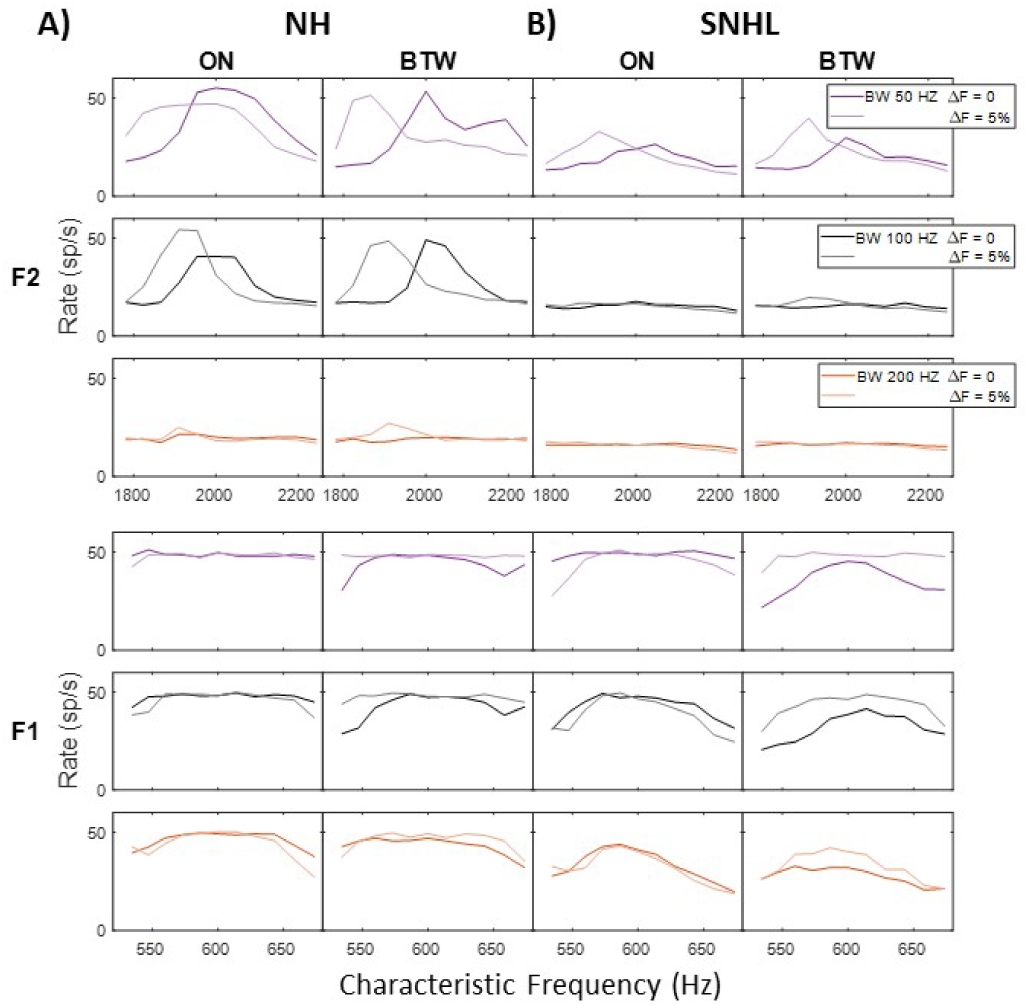
IC BS model rate profiles for inputs with (A) NH, and (B) Elevated thresholds corresponding to 12 dB loss at 600 Hz and 18 dB loss at 2000 Hz (AN model parameters: C_IHC_=0.5, C_OHC_ = 0.5). IC model rate profiles are illustrated for stimuli that differed by ΔF = 5%, which is above discrimination threshold for either NH or listeners with the audiometric thresholds included in (B) (cf., Fig. 5, note log_10_(5) ≈ 0.7). Model rate profiles are shown for the three formant bandwidths using line colors matched to Fig. 3: 50-Hz BW, purple; 100-Hz, black; 200-Hz, orange. For each BW, the darker line corresponds to the standard stimulus, and the lighter line is the response to a formant shifted down by 5% of the standard formant frequency (2000 Hz or 600 Hz). Each IC model profile is for 11 frequency channels spanning 1/3-octave centered at the formant frequency. Model threshold estimates in Fig. 11 were based on the sum of squared errors between rate profiles. The rate profiles illustrated here are based on averages of 10 trials.

Examples of model IC BS population responses to stimulus pairs for different conditions (F1 and F2, ON and BTW) and for different formant bandwidths (colors) are illustrated in Fig. 10. Figure 10A shows NH models responses, and Fig. 10B shows responses for a model that received inputs from an AN model with SNHL (see caption, Fig. 8B). Squared differences between the IC rate profiles for the standard formant frequencies (dark lines, either F1 = 600 H or F2 = 2000 Hz) and for the target formants that were shifted down (light lines, shifted by 5% of the formant frequency in the examples shown) were used as the decision variable to estimate model thresholds. The same 2-pair (i.e., 4-interval) stimuli were used for the model responses for each trial as were used in listening tests, and the pair of model responses with the larger squared rate difference was selected as the target interval. Note that for the model with NH (Fig. 10A), the model IC BS rate profiles in response to F2 (top row) shifted to the left when the formant frequency shifted to a lower frequency (lighter lines), as compared to the standard formant frequency of 2000 Hz (darker lines). The sharper IC BS rate profiles in response to the BTW condition with respect to the ON condition were due to differences in the nature of the stimulus fluctuations when the formant was aligned with a harmonic, as opposed to when it fell between two harmonics (Lyzenga and Horst, 1997; Tan and Carney, 2006; Henry et al., 2017).

The NH model rate profiles in response to F1 (Fig. 10A, bottom row) had relatively small differences over the 1/3-octave range of CFs that were included in the simulations, yet the squared differences between responses across this limited CF range explained listener thresholds (Fig. 11). Interestingly, the differences in NH model rate profiles for the F1 BTW condition were only slightly larger than those for the ON condition, but this greater difference predicted the lower thresholds for the BTW condition that were observed for the youngest listeners with good thresholds (Fig. 11, and Fig. 5, red arrows). Note that the model IC BS rate profiles in response to F1 (Fig. 10, bottom row) do not show simple leftward shifts when the formant frequency was shifted lower (compare light to dark lines for each bandwidth, or color, in Fig. 10). Instead, the reduced IC rates in response to the shifted formants (lighter lines) reflect the larger fluctuations that occur in AN responses when F1 was shifted away from a harmonic (see Fig. 1), and the subsequent suppression of the model IC BS responses by fluctuations on their inputs.

**Figure 11.**
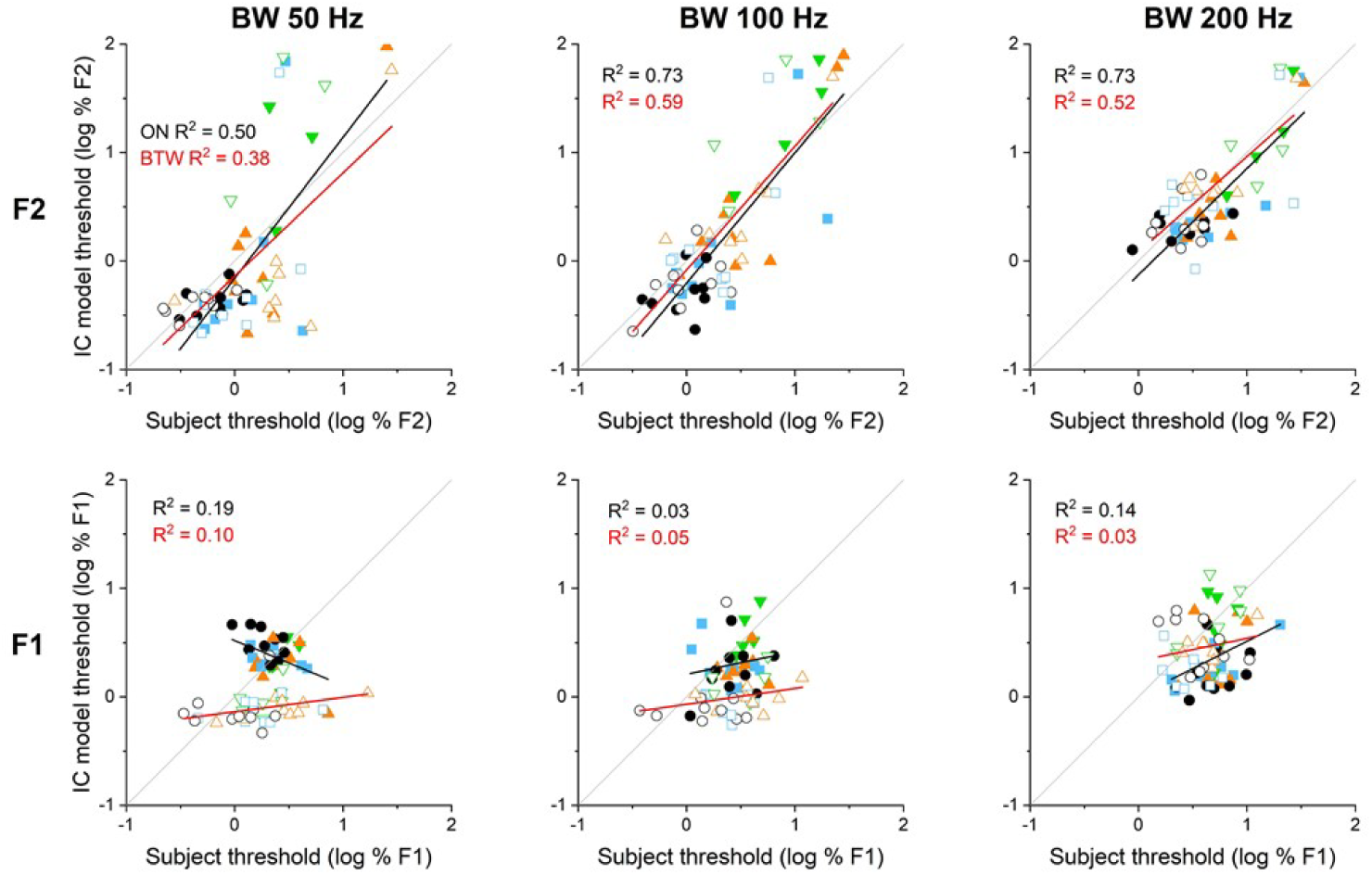
IC-model thresholds as a function of subject thresholds. Symbol types, colors, and lines as in Fig. 9. Compared to the AN model, the IC model better represented ON and BTW human subject data for the F2 condition at all three bandwidths (*cf*. corresponding panels in Fig. 9). For F1, the IC model generally matched the values of the ON subject data at all three bandwidths, but consistently had lower thresholds than the human data for the BTW condition (open symbols, bottom row). Model thresholds were based on 30 repetitions of single-trial responses that were used to estimate logistic functions (see Methods).

The IC model results as a function of subject thresholds are presented in Fig. 11. Compared to the AN model, the IC-model threshold estimates better matched human thresholds for the F2 condition, and were well correlated for the 100- and 200-Hz bandwidth conditions (with correspondingly greater R^2^ regression values; top row, Fig. 11). IC-model F2 threshold estimates were slightly lower than subject thresholds (i.e., regression lines were below the unity line), suggesting that the IC model was sufficient to explain most listeners’ F2 thresholds, whereas the AN-model thresholds were not (Fig. 9). The IC model also performed better than the AN model in estimating the subjects’ F1 thresholds (cf. linear regressions in the bottom rows of Figs. 9 and 11). For the F1 ON condition the IC model correctly predicted the listeners’ threshold values, whereas for the F1 BTW condition, with the exception of subjects with the lowest thresholds, IC-model thresholds were substantially lower than the corresponding listeners’ thresholds (i.e., points and red regression lines fell below the gray unity line in Fig. 11, bottom row), a trend that decreased with increasing bandwidth. These results indicate that detection cues available to the IC model might also be used by human subjects, especially for the F2 condition at 100 and 200 Hz (both ON and BTW). For the F1 condition, the model apparently takes advantage of enhanced fluctuation cues present in the BTW condition, but only the most sensitive human subjects apparently used those cues.

The model simulations were unable to estimate thresholds for F2 discrimination for the listeners with the most hearing loss. Specifically, the AN model rate profiles were not successful in estimating F2-discrimination thresholds for listeners with audiometric thresholds greater than 30 dB at 2 kHz (8/34 listeners). The IC model, which took advantage of not only rate but temporal features in AN responses, was more successful and only failed to estimate F2-discrimination thresholds for listeners with greater than 40 dB HL at 2 kHz (5/34 listeners.) Although not tested here, it is possible that a model based on a larger number and/or a wider range of CFs could predict discrimination thresholds for the listeners with greatest hearing loss. However, it is also likely that the standard assumption used to incorporate hearing loss in the peripheral model, which attributed 2/3 of hearing loss to OHC dysfunction and 1/3 to reduced sensitivity of IHCs, may not apply to listeners with the largest audiometric threshold elevations.

## DISCUSSION

The goal of this study was to test the hypothesis that NF cues support encoding of formant frequencies in vowels. Two factors that affect NF coding in computational models of midbrain responses to vowels are formant bandwidth and SNHL. The results of manipulating the bandwidth of formants in synthetic vowel-like sounds and by testing listeners with a range of SNHL generally supported the hypothesis. Furthermore, these results showed a surprising *insensitivity* of F1 discrimination to SNHL and a surprising *sensitivity* of F2 discrimination to SNHL. The NF-based IC-level model was largely able to capture the trends in these results, although the IC model predicted lower thresholds for most listeners to the BTW condition. The BTW condition has been explored in previous studies of NH subjects (humans: Lyzenga and Horst, 1995; budgerigars: Henry et al., 2017). Model thresholds were consistent with those of young listeners with NH who perform better in the BTW condition as compared to the ON condition, as expected due to stronger fluctuation cues in response to BTW stimuli (Tan and Carney, 2006). However, most other listeners in our test group, especially those who were not in the youngest group or had any measurable elevation of audiometric thresholds, did not seem to take advantage of these fluctuation cues, based on the fact that they did not have improved performance for the BTW condition.

IC-model thresholds were particularly successful at predicting the strong dependence of F2-discrimination thresholds on SNHL (Fig. 11). In contrast, a model based on AN rate profiles did not predict the trends in threshold with SNHL and had thresholds that were generally elevated with respect to listeners’ thresholds (Fig. 9). The improved predictions of thresholds by the IC-model rate profile, as compared to the AN model rate profile, suggests that NFs play an important role in coding of the vowel spectra. AN rate profiles provide a representation of the “excitation pattern” associated with the responses of the inner ear, but do not include the temporal patterns of the responses. In contrast, the IC-model rate profile combines the representations of energy and low-frequency temporal fluctuations. In this study SNHL was included only in the peripheral model. SNHL increases the NFs in peripheral responses due to reduced capture of channels tuned near spectral peaks, an effect observed in physiological studies of the AN following noise-induced SNHL (e.g., Kale and Heinz, 2012). It is possible that sensorineural hearing loss also affects sensitivity to NFS at the level of the midbrain. Our model for modulation tuning (Nelson and Carney, 2004) would predict broadening of MTFs as a result of reduced inhibition in IC inputs. There are currently no direct reports of SNHL effects on the bandwidths of IC MTFs, but the balance of excitation and inhibition in the IC is affected by SNHL (Barsz et al., 2007), age (Caspary et al., 1999), acoustic trauma (Mulders and Robertson, 2013; Ma et al., 2020), and cochlear ablation (Vale and Sanes, 2002; Vale et al., 2004). There are also reports of changes in the distribution of MTF types following noise-induced trauma (Heeringa and van Dijk, 2019) and in aged animals (Walton et al., 2002), although a study of the effects of SNHL on temporal coding in the mouse IC reported no significant effect on neural thresholds for temporal gap detection (Walton et al., 2008). Despite the current lack of direct physiological evidence to support modification of model MTF bandwidths, one can assume that the increase in thresholds for listeners with SNHL is explained, at least in part, by an increase in bandwidth of MTFs that would reduce the contrast of neural representation of formants. Reductions in the contrast of NFs at the level of peripheral and in the central neural populations would both tend to elevate thresholds in listeners with SNHL.

In addition to analyzing and modeling perceptual thresholds for F1 and F2 discrimination, we compared thresholds to listeners’ performance on a speech-in-noise task. QSIN scores were correlated to thresholds for F2, most strongly for the F2 BTW condition (Fig. 7, Table 1). Thus, this condition may provide an alternative to intelligibility testing for quantifying sensitivity of individual listeners to changes in speech sounds. A potential benefit of using the formant-frequency discrimination task to predict speech intelligibility is that it likely to be less sensitive to cognitive factors associated with speech testing, including a native English-speaking background. This result is consistent with earlier studies, based on vowel similarity testing, which reported that F2 discrimination is an important factor in multiple-regression analyses relating speech perception to psychophysical thresholds in listeners with SNHL, and that F1 is weighted more strongly in listeners with SNHL (Dreschler and Plomp, 1980, 1985).

Future studies will explore strategies for enhancing intelligibility by manipulating neural-fluctuation profiles. The potential implication of elevated DL_FF_s for F2 on intelligibility is illustrated by the loss of resolution on the F2 axis of the English-language vowel space (Fig. 12). For our listeners, F1 DL_FF_s were not significantly elevated by HL at 500 Hz, thus one would expect that their resolution along the F1 axis of a 2D vowel space would not be affected. However, F2 DL_FF_s were significantly elevated by HL at 2 kHz, by as much as 35% for the listeners in this study with the largest SNHL. Figure 11 shows the Peterson and Barney (1952) vowel space for English to illustrate the effect of a 35% decrease in F2 resolution, i.e., a 35% increase in the size of vowel clusters. Considerable overlap between vowel clusters is consistent with elevated QSIN scores for listeners with high F2 DL_FF_s. The increase in overlap of spaces associated with vowel contrasts depending on F2 is generally consistent with vowel confusions in listeners with SNHL reported by Van Tasell et al. (1987) (e.g., the most common confusion reported was between /ʌ/ and /a/, with other common confusions being /ʌ/ and /æ/, /ɛ/ and /æ/, and /a/ and /æ/).

**Figure 12.**
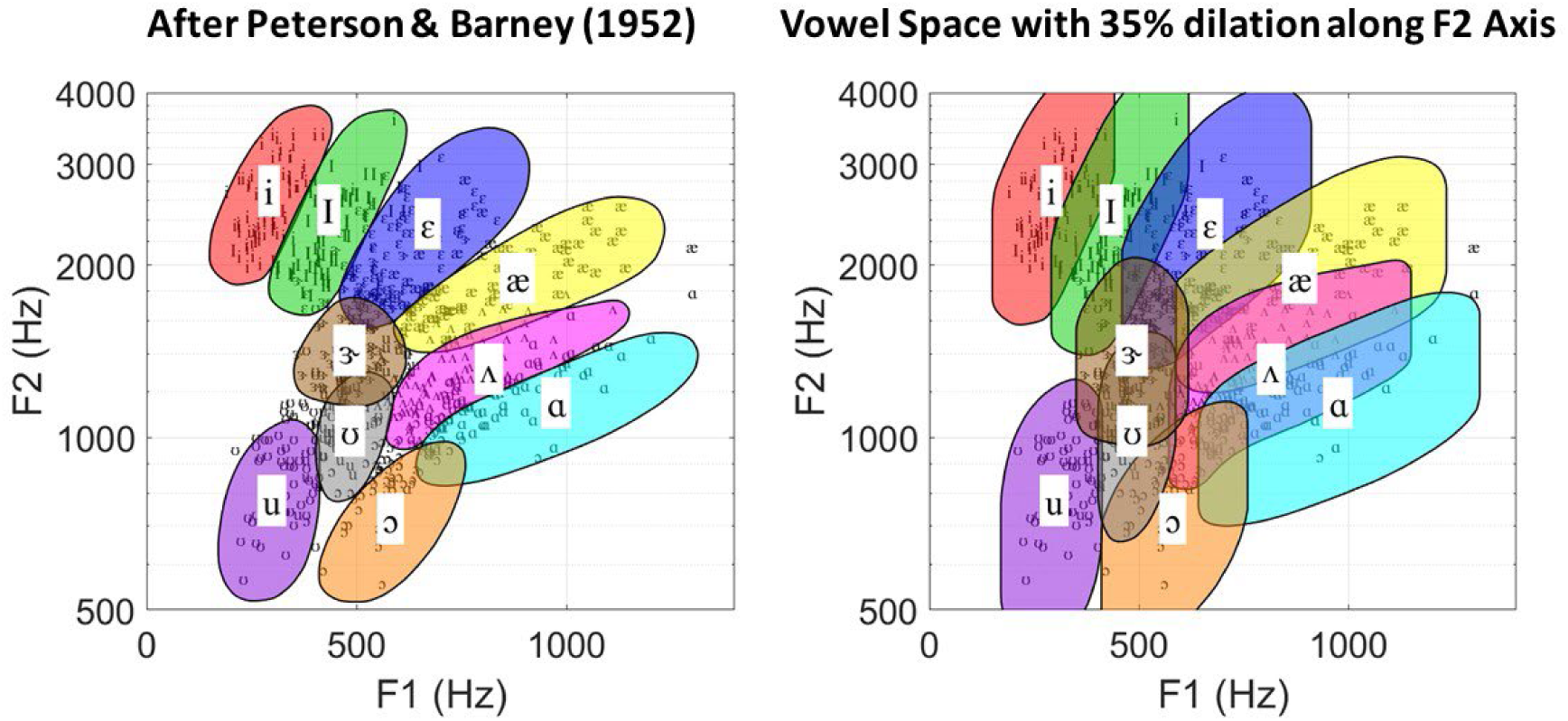
Vowel space (A) with ‘dilation’ corresponding to 35% increase in F2 range (±17% with respect to Peterson and Barney, 1952) based on changes in DL_FF_s for listeners with SNHL. Schematic visualization of the implications of elevated FFDLs for vowel discrimination. NOTE: Threshold for listeners with most hearing loss in F2, BW=100 Hz condition was 35% [log10(35%) = 1.54 on log axis] for both ON and BTW conditions. This listener required a 35% difference in the frequency of F2 to reliably detect a change, in comparison to subjects with 0 dB HL who required ∼1% difference. Diagram shows schematically the increase in overlap of the vowels in the vowel space that might correspond to higher DL_FF_s for F2.

Our results shed new light on the structure of vowel contrasts and systems. In a well-documented, cross-linguistic phenomenon, observed vowel contrasts pattern in consistent ways, dispersing themselves within a space defined by the acoustic parameters of the vowels’ first two formants, F1 and F2, referred to as Dispersion Theory (Liljencrants and Lindblom, 1972; Lindblom 1986; Schwartz et. al. 1997a,b; Becker-Kristal, 2010). A given vowel space is defined by the highest front vowel [i] and the lowest vowel [a] (Becker-Kristal, 2010), with larger systems occupying a larger space. However, the larger vowel systems differ from smaller systems in the number of vowel contrasts, mainly along the F1 (height) parameter. Attempts at modeling the vowel space overpredict the number of vowel contrasts that appear along the F2 (backness) parameter (Liljencrants and Lindblom, 1972; Schwartz et al., 1997a). The acoustics of vowels sounds cannot explain these vowel-dispersion patterns, thus nearly every study in this area has called on the need to understand the constraints of the auditory system on vowel patterns. Our finding that formant-frequency discrimination in the F2 dimension is particularly vulnerable to hearing loss, whereas discrimination in the F1 dimension is not, is consistent with the relative importance of F1 in vowel contrasts in most languages. Additionally, the spectral magnitude of F2 is inherently lower than that of F1 (Klatt, 1980), exacerbating the effect of hearing loss in the F2 frequency region.

## ACKNOWLEDGEMENT

Supported by NIH-R01-DC001641

## REFERENCES

Barsz, K., Wilson, W. W., & Walton, J. P. (2007). Reorganization of receptive fields following hearing loss in inferior colliculus neurons. Neuroscience, 147(2), 532–545.

Becker-Kristal, Roy (2010) Acoustic Typology of Vowel Inventories and Dispersion Theory: Insights from a Large Cross-Linguistic Corpus Ph.D. Dissertation, University of California, Los Angeles

Carney, LH (2018) Supra-threshold hearing and fluctuation contrast: Implications for sensorineural and hidden hearing loss, JARO 19:331–352.

Carney, LH, Li, T, McDonough, JM (2015) Speech Coding in the Brain: Representation of Formants by Midbrain Neurons Tuned to Sound Fluctuations. eNeuro 2(4), 1–12. e0004-15.2015 1–1. (DOI: 10.1523/ENEURO.0004-15.2015).

Carney, LH, Kim DO, Kuwada, S (2016) Speech Coding in the Midbrain: Effects of Sensorineural Hearing Loss, in P. van Dijk, D. Başkent, E. Gaudrain, E. de Kleine, A. Wagner, C. Lanting (Eds.) Physiology, Psychoacoustics and Cognition in Normal and Impaired Hearing,. Springer. Advances in experimental medicine and biology. 894:427–35.

Carney, LH, and JM McDonough (2019) Nonlinear auditory models yield new insights into representations of vowels, Atten Percept Psychophys, 81(4):1034–1046. DOI: 10.3758/s13414-018-01644-w.

Caspary, D. M., Holder, T. M., Hughes, L. F., Milbrandt, J. C., McKernan, R. M., & Naritoku, D. K. (1999). Age-related changes in GABAA receptor subunit composition and function in rat auditory system. Neuroscience, 93(1), 307–312.

Delgutte B, Kiang NYS (1984). Speech coding in the auditory nerve: I. Vowel-like sounds. J Acoust Soc Am 75: 866–878.

Deng, L., & Geisler, CD (1987). A composite auditory model for processing speech sounds. The Journal of the Acoustical Society of America, 82(6), 2001–2012.

Dreschler, W. A., & Plomp, R. (1980). Relation between psychophysical data and speech perception for hearing-impaired subjects. I. The Journal of the Acoustical Society of America, 68(6), 1608–1615.

Dreschler, W. A., & Plomp, R. (1985). Relations between psychophysical data and speech perception for hearing-impaired subjects. II. The Journal of the Acoustical Society of America, 78(4), 1261–1270.

Hamza, Y., Farhadi, A., Schwarz, D.M., McDonough, J.M., Carney, L.H. (2022, in review) Representations of fricatives in sub-cortical model responses: comparisons with human perception, BioRxiv. 2022.10.24.513605; doi: https://doi.org/10.1101/2022.10.24.513605.

Heeringa, A. N., & van Dijk, P. (2019). Neural coding of the sound envelope is changed in the inferior colliculus immediately following acoustic trauma. European Journal of Neuroscience, 49(10), 1220–1232.

Henry, K.S., Amburgey, K.N., Abrams, K.S., Idrobo, F., Carney, L.H. (2017) Formant-frequency discrimination of synthesized vowels in budgerigars (*Melopsittacus undulatus*) and humans, JASA. 142(4), 2073–2083.

Kale, S., & Heinz, M. G. (2012). Temporal modulation transfer functions measured from auditory-nerve responses following sensorineural hearing loss. Hearing research, 286(1-2), 64–75.

Killion, M. C., Niquette, P. A., Gudmundsen, G. I., Revit, L. J., & Banerjee, S. (2004). Development of a quick speech-in-noise test for measuring signal-to-noise ratio loss in normal-hearing and hearing-impaired listeners. The Journal of the Acoustical Society of America, 116(4), 2395–2405.

Kim, D. O., Carney, L., & Kuwada, S. (2020). Amplitude modulation transfer functions reveal opposing populations within both the inferior colliculus and medial geniculate body. Journal of Neurophysiology, 124(4), 1198–1215.

Klatt, D. H. (1980). Software for a cascade/parallel formant synthesizer. Journal of the Acoustical Society of America, 67(3), 971–995.

Krishna, B. S., & Semple, M. N. (2000). Auditory temporal processing: responses to sinusoidally amplitude-modulated tones in the inferior colliculus. Journal of neurophysiology, 84(1), 255–273.

Nelson, P. C., & Carney, L. H. (2007). Neural rate and timing cues for detection and discrimination of amplitude-modulated tones in the awake rabbit inferior colliculus. Journal of neurophysiology, 97(1), 522–539.

Joris, P. X., Schreiner, C. E., & Rees, A. (2004). Neural processing of amplitude-modulated sounds. Physiological reviews, 84(2), 541–577.

Lindblom, B. (1986). Phonetic universals in vowel systems. In: Ohala, J., Jaeger, J. (eds.) Experimental Phonology. Orlando: Academic Press, 13–44.

Liljencrants, J., & Lindblom, B. (1972). Numerical simulation of vowel quality systems: The role of perceptual contrast. Language, 48, 839–862. doi:10.2307/411991

Lyzenga, J., & Horst, J. W. (1995). Frequency discrimination of bandlimited harmonic complexes related to vowel formants. The Journal of the Acoustical Society of America, 98(4), 1943–1955.

Lyzenga, J., & Horst, J. W. (1997). Frequency discrimination of stylized synthetic vowels with a single formant. The Journal of the Acoustical Society of America, 102(3), 1755–1767.

Lyzenga, J., & Horst, J. W. (1998). Frequency discrimination of stylized synthetic vowels with two formants. The Journal of the Acoustical Society of America, 104(5), 2956–2966.

Ma, L., Ono, M., Qin, L., & Kato, N. (2020). Acoustic trauma induced the alteration of the activity balance of excitatory and inhibitory neurons in the inferior colliculus of mice. Hearing Research, 391, 107957.

Mulders, W. H. A. M., & Robertson, D. (2013). Development of hyperactivity after acoustic trauma in the guinea pig inferior colliculus. Hearing research, 298, 104–108.

Peterson, G. E., & Barney, H. L. (1952). Control methods used in a study of the vowels. The Journal of the acoustical society of America, 24(2), 175–184.

Rosenberg, A. E. (1971). Effect of glottal pulse shape on the quality of natural vowels. The Journal of the Acoustical Society of America, 49(2B), 583–590.

Schwartz, J.L., Boë, L.J., Vallée, N., Abry, C. (1997a). Major trends in vowel system inventories. J. Phonetics 25: 233–253

Schwartz, J.L., Boë, L.J., Vallée, N., Abry, C. (1997b). The Dispersion-Focalization Theory of vowel systems. J. Phonetics 25:255–286

Tan and Carney (2006), Predictions of Formant-Frequency Discrimination in Noise Based on Model Auditory-Nerve Responses, J. Acoust. Soc. Am. 120:1435–1445. PMCID: PMC2572872.

Young, E. D., & Sachs, M. B. (1979). Representation of steady-state vowels in the temporal aspects of the discharge patterns of populations of auditory-nerve fibers. The Journal of the Acoustical Society of America, 66(5), 1381–1403.

Vale, C., Juíz, J. M., Moore, D. R., & Sanes, D. H. (2004). Unilateral cochlear ablation produces greater loss of inhibition in the contralateral inferior colliculus. European Journal of Neuroscience, 20(8), 2133–2140.

Vale, C., & Sanes, D. H. (2002). The effect of bilateral deafness on excitatory and inhibitory synaptic strength in the inferior colliculus. European Journal of Neuroscience, 16(12), 2394–2404.

Van Tasell, D. J., Fabry, D. A., & Thibodeau, L. M. (1987). Vowel identification and vowel masking patterns of hearing-impaired subjects. The Journal of the Acoustical Society of America, 81(5), 1586–1597.

Walton, J. P., Barsz, K., & Wilson, W. W. (2008). Sensorineural hearing loss and neural correlates of temporal acuity in the inferior colliculus of the C57BL/6 mouse. Journal of the Association for Research in Otolaryngology, 9, 90–101.

Walton, J. P., Simon, H., & Frisina, R. D. (2002). Age-related alterations in the neural coding of envelope periodicities. Journal of neurophysiology, 88(2), 565–578.

Zaar, J., & Carney, L. H. (2022). Predicting speech intelligibility in hearing-impaired listeners using a physiologically inspired auditory model. Hearing Research, 108553.

Zilany, M. S., & Bruce, I. C. (2007). Representation of the vowel/ε/in normal and impaired auditory nerve fibers: model predictions of responses in cats. The Journal of the Acoustical Society of America, 122(1), 402–417.

Zilany, M.S.A., Bruce, I.C., and L. H. Carney (2014) Updated parameters and expanded simulation options for a model of the auditory periphery. J Acoust Soc Am 135:283–286.

